# Culturally Universal Temporal Dynamics of Statistical Learning in Children’s Song Contributes to Phase Entrainment and Production of Novel Information

**DOI:** 10.1101/2023.07.17.549433

**Authors:** Tatsuya Daikoku

## Abstract

Statistical learning is intimately linked to brain development. For example, statistical learning of language and music starts at an early age and is shown to play a significant role in acquiring the delta-band rhythm that is essential for language and music learning. However, it remains unclear how auditory cultural differences affect the statistical learning process and the resulting probabilistic and acoustic knowledge acquired through it.

This study examined how children’s songs are acquired through statistical learning. This study used a Hierarchical Bayesian Statistical Learning (HBSL) model, mimicking the statistical learning processes of the brain. Using this model, we conducted a simulation experiment to visualize the temporal dynamics of perception and production processes through statistical learning among different cultures. The model learns the children’s songs (MIDI files) including English, German, Spanish, Japanese, and Korean songs as the training data. This study investigated how the model transforms the internal model over 15 trials of learning in each song. Furthermore, using the probability distribution of each model over 15 trials of learning each song, new songs were probabilistically generated.

The results suggest that, in learning processes, chunking and hierarchical knowledge increased gradually through 15 rounds of statistical learning for each piece of children’s songs. In production processes, statistical learning leads to the gradual increase of delta-band rhythm (1-3 Hz). Furthermore, by combining the acquired chunks and hierarchy through statistical learning, new music can be generated gradually in comparison to the original songs (i.e., the training songs). These findings were observed consistently, irrespective of culture.

The present study indicates that the innate statistical learning capacity of the brain, irrespective of cultural background, contributes to the acquisition and generation of delta-band rhythm, which is critical for acquiring language and music. It is suggested that cultural differences may not significantly modulate the statistical learning effects since statistical learning and slower rhythm processing is both essential functions in the human brain across cultures. Furthermore, statistical learning of children’s songs leads to the acquisition of hierarchical knowledge and the ability to generate novel music. This study provides a novel perspective on the developmental origins of creativity and the importance of statistical learning through early development.

## 1. Introduction

### 2.1. Statistical Learning and Emergence of Individuality

Music is a ubiquitous element in all cultures and is instrumental in shaping the unique characteristics that define each one [1]. Through the learning of music, humans develop culturally specific or universal knowledge. Although music plays a crucial role in human development, the underlying mechanisms that enable the brain to learn and produce music remain poorly understood.

In recent years, a growing body of studies has tried to explain the learning and production mechanisms of music based on the general principle of predictive processing in the brain [2]. Predictive processing of music works to minimize the prediction error (i.e., surpirse) between the bottom-up auditory signals including melody, rhythm, and musical chords and the top-down predictive signals based on internal music models of the brain [3,4].

Within the framework of the brain’s predictive processing, it has been proposed that the acquisition of musical knowledge may be underpinned by statistical learning mechanisms [5,6]. The statistical learning is an innate function that is closely linked to brain development [7] and contributes to the perception and production of music and language. The basic mechanism involves calculating the statistical probability of environmental information (particularly the transition probability of sequential information) and the uncertainty of probability distribution, and predicting future information based on an internal probabilistic model acquired through statistical learning.

Importantly, the uncertainty and probability are not universally inherent in music per se but is instead shaped by an amount of individual’s auditory experiences. For example, when non-native individuals listen to culturally specific ethic music, the uncertainty in predicting the probable subsequent sound is typically high, making prediction more difficult. In contrast, people within the ethnic group who regularly listen to such music often find it easier to predict sounds (i.e., less surprise) due to their lower level of uncertainty. This is a result of individuals constantly updating their internal models through extended periods of statistical learning, thereby generating an appropriate music probability model within the culture. Neural and computational studies have suggested that the individual traits related with music “production” (i.e., composition) [8-10] and perception [11-13] are associated with past experiences in statistical learning.

### 2.2. Reliability of Prediction

Auditory experience affects not only uncertainty, prediction, and surprise but also the “reliability” of predictions. For example, the degree of reliability varies even when the transition probability is identical at 90% between the case of experiencing A-to-B transitions nine times out of 10 trials and the case of experiencing them 90 times out of 100 trials. Additionally, the brain can rely more on events with low transition probability if they have been experienced 10 out of 100 times instead of only once out of 10. Intuitively, one is in a state of recognizing that this event is “reliably” unpredictable.

Neurophysiological studies [14,15] have demonstrated a progressive representation of statistical learning effects with an increase in the number of learning repetitions. This indicates that the brain’s statistical learning is based on Bayesian inference, which enhances the reliability of probabilities with experience, as opposed to maximum likelihood estimation, which does not vary with learning repetitions. However, most studies of statistical learning have referred to maximum likelihood estimation based on Markov models or n-gram models that do not consider the “reliability” of probabilities, and thus, have not considered the effect of learning repetitions.

To resolve such problems of these statistical learning studies, this study developed a computational model of the brain’s statistical learning, referred to as “Hierarchical Bayesian Statistical Learning (HBSL)” incorporating the Bayesian reliability of probabilities into a Markov model. We then used this model to examine the learning process when lots of music are repetitively learned.

### 2.3. Hierarchically Structured Building and Its Phase Entrainment

Statistical learning essentially contributes to chunking, which involves identifying information units with high transition probabilities from sequential information such as short phrases and words. Prior research has investigated the neural and computational mechanisms underlying chunking through statistical learning. In contrast, recent studies have suggested two types of *“hierarchical”* statistical learning systems [10,16]. The first system constitutes the fundamental function of statistical learning, which groups chunks of information with high transition probabilities and integrates them into a cohesive unit. The second system involves statistical learning that arranges various chunked units to form a hierarchical syntactic structure (Figure 1). Therefore, statistical learning plays a critical role in acquiring the hierarchy, a unique and essential feature of language and music [17].

**Figure 1.**
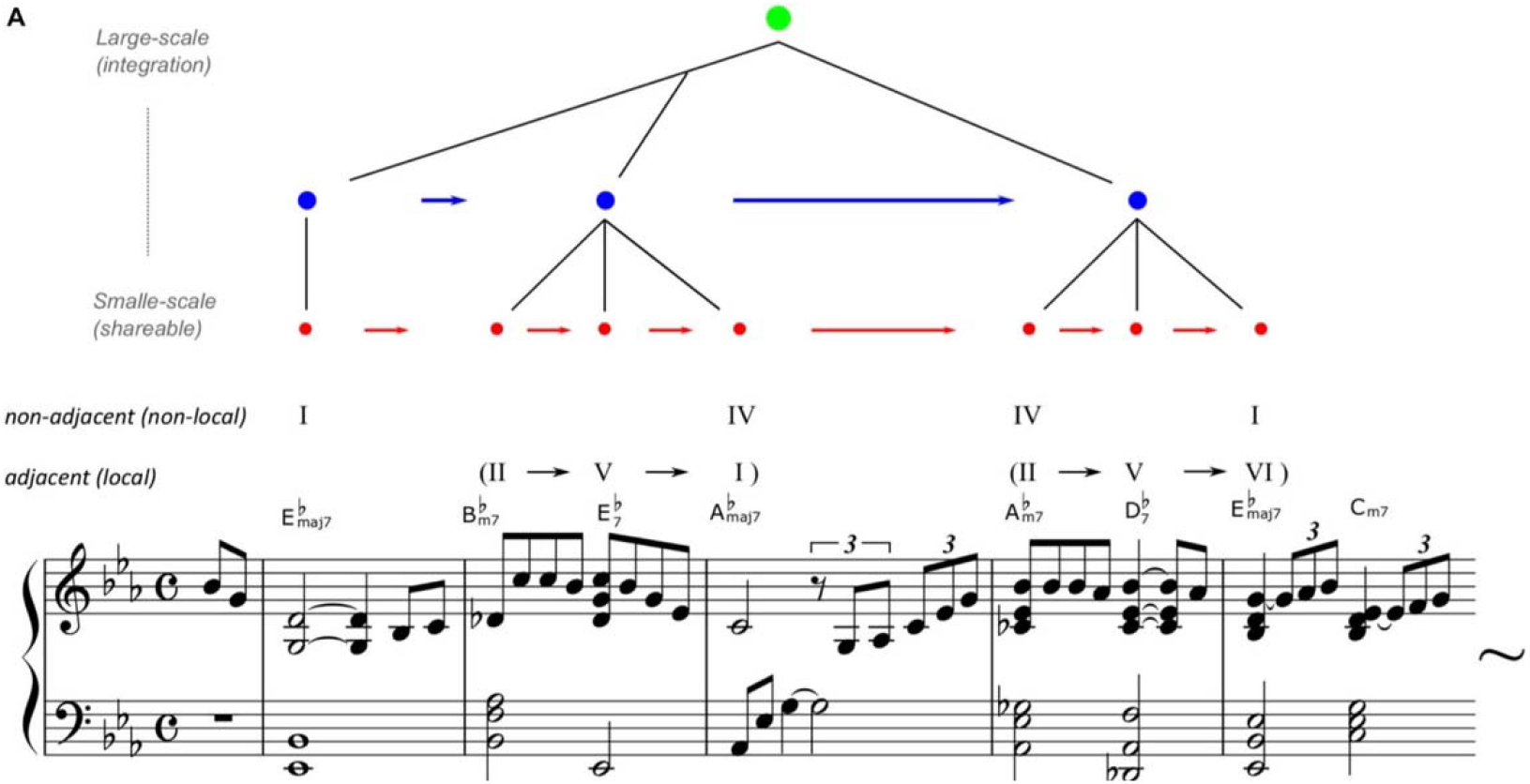
An example of hierarchical statistical learning of music. Reprinted from Daikoku et al. (2021) [10]. Misty by Errol Garner, composed in 1954. The arrangement, chord names, and symbols are simplified (just major/minor, flat, and 7th note) to account for the two-five-one (II–V(7)–I) progression. For example, jazz music has general regularities in chord sequences such as the so-called “two-five-one (II–V–I) progression.” Such syntactic progression frequently occurs in music, and therefore, the statistics of the sequential information have high transitional probability and low uncertainty. Thus, once a person has learned the statistical characteristics, it can be chunked as a commonly used unit among improvisers. In contrast, the ways of combining the chunked units (e.g., combining among different blue circled units) are different between musicians.

Particularly, the hierarchical structure of rhythms is essential for the acquisition of music and language [18]. The lower hierarchy corresponds to a frequency band of syllable and musical notes (e.g., quarter notes) around 4-12Hz while the higher hierarchy corresponds to prosody, intonation, and long musical note (e.g., half notes) around 1-3Hz [19]. There are also rhythms around 12-30Hz that correspond to phonemes or sound onsets at even lower levels of the hierarchy.

Such a hierarchical structure of rhythm can be visualized from the amplitude modulation (AM) envelope of sound waveforms [20,21]. Evidence has shown that neural oscillation in the human brain phase synchronizes the AM hierarchy of auditory rhythm [22]. Such a synchronization or phase entrainment contributes to parsing of the sound signal into each unit of the hierarchy [23]. Further, even in the absence of acoustic (e.g., AM) cues to hierarchical structure, it has been demonstrated that the brain is phase entrained to the hierarchical rhythm simply from the syntactic but not acoustic structure [24]. This raises the possibility that, even if the input information has a weak acoustic hierarchy, the brain can produce output information with an acoustic hierarchical structure by syntactically processing the hierarchical structure in the brain.

A study comparing the hierarchical rhythmic structure across various types of sounds (nature, speech, instrumental music, song, animal sounds, etc.) has demonstrated that the slower band hierarchy (i.e., 1-3 Hz) is especially pronounced in children’s songs and speech directed towards infants and children (referred to as infant/child-directed speech) compared to other types of sounds [19] and possibly regardless of culture [25]. Further, evidence has shown that infant is first phase-entrained to the slower rhythm (1-3Hz) [26], suggesting that such a slower-band rhythm is important for early learning of language and music. Thus, children’s songs and child-directed speech may emphasize 1-3 Hz rhythms to facilitate the acquisition of rhythm hierarchy in early auditory learning. Alternatively, it could be hypothesized that children’s songs emphasize 1-3 Hz rhythms because they syntactically and acoustically learn these rhythms first.

Evidence has shown that the ability of 1-3 Hz phase entrainment is closely linked to statistical learning capacity [27]. By statistical learning, the neural oscillations can be entrained to the statistically chunked information that has slower rhythm (e.g., 1Hz word rhythm that has three 3Hz syllables, [28,29]). Thus, statistical learning plays a critical role in phase entrainment of the slower rhythm around 1-3Hz, essential for early learning of of music. However, it is unclear how music stimuli are chunked into a 1-3Hz unit and how they are manifested as 1-3 Hz rhythms when producing a new musical information through statistical learning, and whether different internal models that may vary with culture and experience affect the processing of 1-3 Hz rhythms.

### 2.4. Purpose of The Present Study

The present study examined the temporal dynamics of perception and production processes through statistical learning, using the HBSL model that considers both reliability and hierarchy, mimicking the statistical learning processes of the brains. The model learns the children’s song (MIDI files) including English, German, Spanish, Japanese, and Korean songs as the training data. This study investigated how the model transforms the internal model over 15 trials of learning in each song. Furthermore, using the probability distribution of these models, new songs were probabilistically generated for each model.

This study hypothesizes that statistical learning leads to gradual increase of 1-3 Hz rhythms in music production. Further, since statistical learning and slower rhythm processing are both essential function in the human brain across culture, it is hypothesized that cultural differences do not significantly modulate the statistical learning effects.

## 2. Results

### 2.1. Learning Process

The total number of chunks and hierarchy generated during statistical learning was measured in each of 15 trials. All results were shown in an external source (https://osf.io/cqmz8/?view_only=b95d94626a364700adb9e1e94384525d). The results revealed that over the course of 15 statistical learning trials, both the number of chunks (as demonstrated by the top left quadrant of Figure 2) and hierarchy (as demonstrated by the right quadrant of Figure 2) gradually increased. Further, the total Bayesian surprise (or total prediction errors) that occurred in each statistical learning was measured by the Kullback-Leibler divergence between a probability distribution P(x) before exposure to a stimulus and a probability distribution after the exposure to a stimulus. The results showed that the Bayesian surprise showed a gradual decrease over the course of 15 statistical learning trials, as demonstrated by the data located in the bottom left quadrant of Figure 2. Finally, these findings were consistent across different cultural groups, indicating the universal phenomena across cultures.

**Figure 2.**
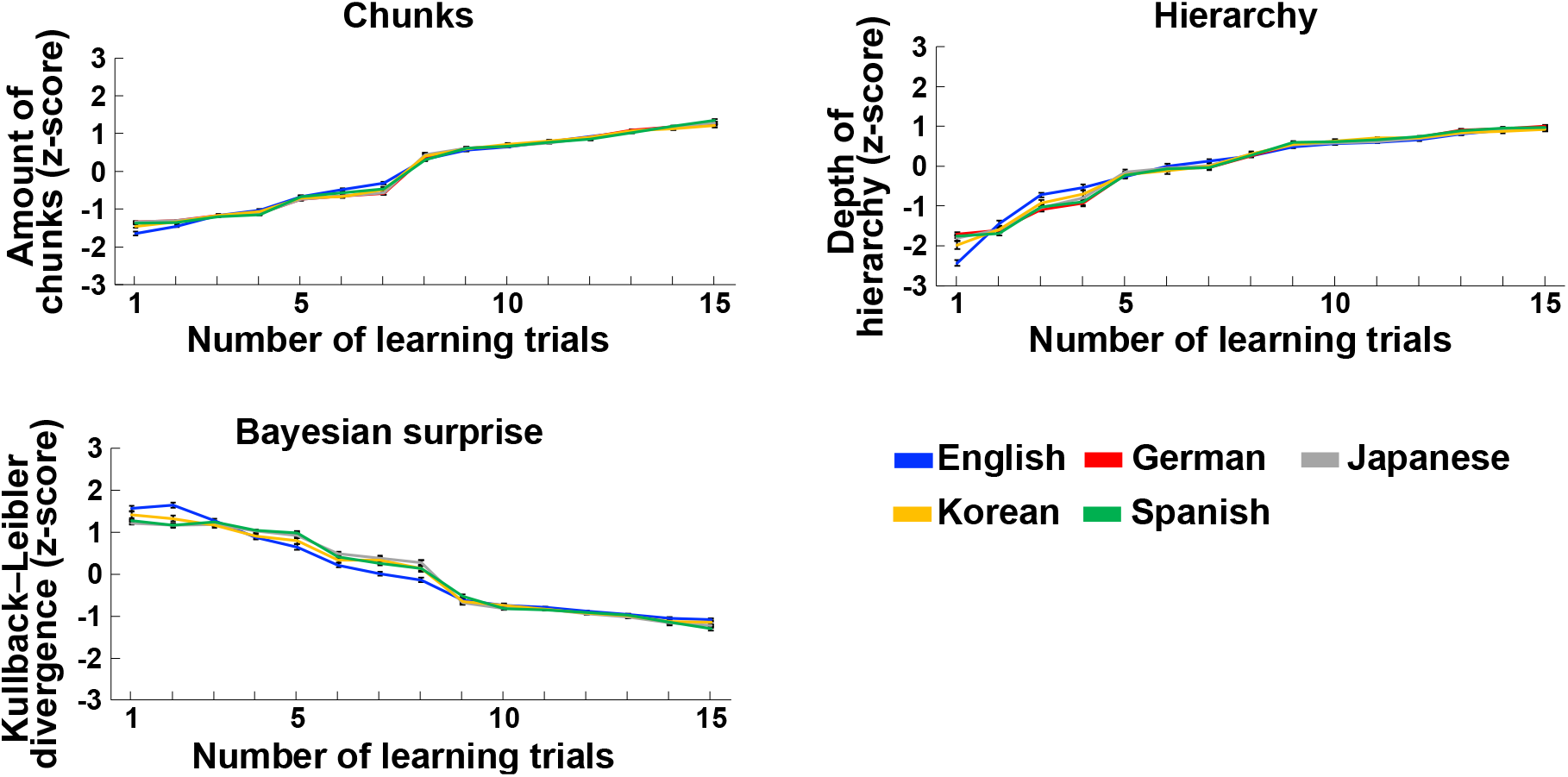
Statistical learning processes in each trial of learning. The top left is the numbers of chunks after statistical learning, the bottom left is the total Bayesian surprise (or prediction error) in learning a piece of music, and the right is the numbers (or depth) of hierarchy after each trial of statistical learning. Blue, red, grey, yellow and green represent children’s songs in English, German, Japanese, Korean and Spanish, respectively. Each value was normalized to z score. The error bars indicate standard error of the mean.

### 2.2. Production Process

#### 2.2.1. Acoustic Dimension

Using the songs composed after each statistical learning, the acoustic property (amplitude modulation waveforms) below 15 Hz, which corresponds to auditory rhythm, were extracted using the Bayesian model called PAD [20]. All results were shown in an external source (https://osf.io/cqmz8/?view_only=b95d94626a364700adb9e1e94384525d). The modulators were converted into time-frequency domains using scalogram. Then, the average frequency power among the 20 songs generated by each model were calculated. The results showed that the composed songs gradually increased lower-band rhythm (1-3 Hz, 1 or 2Hz peak) and decreased higher-band rhythm (3-5Hz, 4Hz peak) (Figure 3). Furthermore, over the course of 15 statistical learning trials, new music was generated gradually in comparison to the original songs (i.e., the training songs). These findings were observed consistently, irrespective of culture. These findings were observed consistently, irrespective of culture.

**Figure 3.**
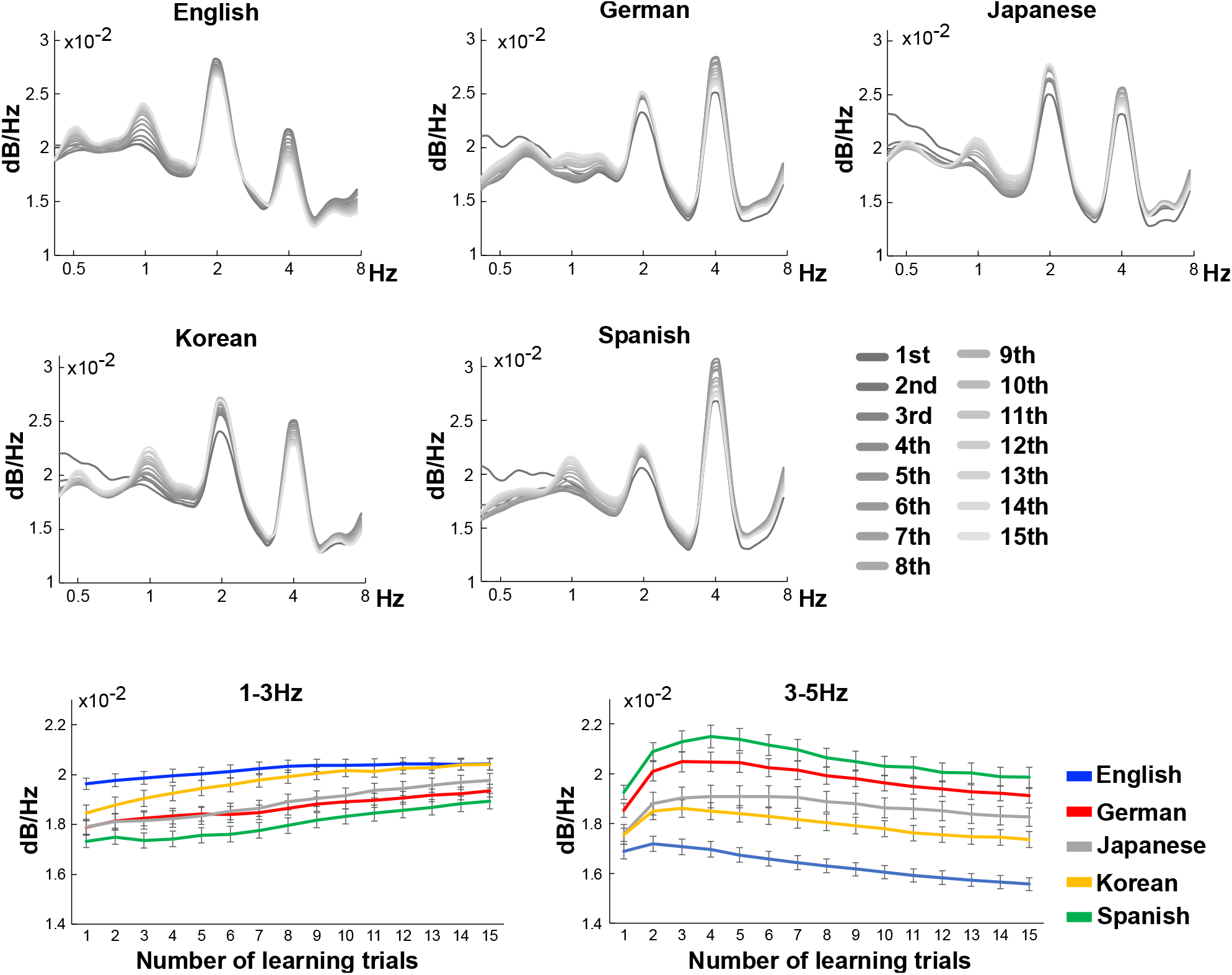
Acoustic properties of the composed music after each trial of statistical learning. The rhythm waveforms (amplitude modulation waveforms) below 15 Hz were extracted using the Bayesian model called PAD [20]). Then the modulators were converted into time-frequency domains using scalogram. Finally, the average frequency power among the 20 songs generated by each model were calculated. Blue, red, grey, yellow and green represent children’s songs in English, German, Japanese, Korean and Spanish, respectively.

#### 2.2.2. Probabilistic Dimension

The average probability distribution of the 20 songs generated by a model after each trial of statistical learning was calculated. The statistical similarity of the probability distributions of music composed after each of the 15 trials of statistical learning was compared to the training data (i.e., original data) using tSNE. The results showed that novel music was generated gradually in comparison to the original songs (i.e., the training songs) over the course of 15 statistical learning trials (Figure 4). These findings were observed consistently, irrespective of culture.

**Figure 4.**
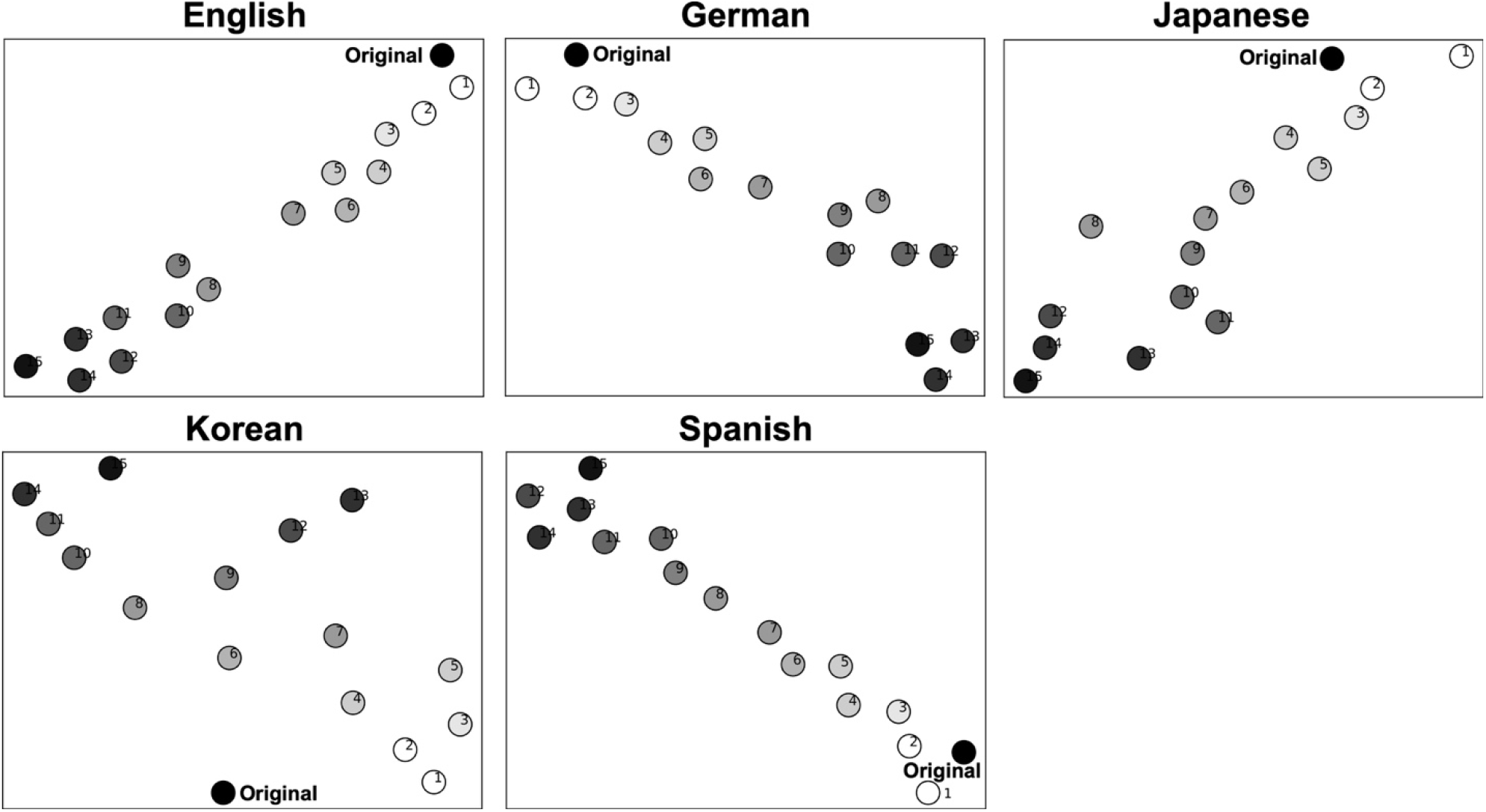
Statistical properties of the composed music after each trial of statistical learning. The average probability distribution of the 20 songs generated by a model after each trial of statistical learning was calculated. The statistical similarity of the probability distributions of music composed after each of the 15 trials of statistical learning was compared to the training data (i.e., original data) using t-distributed stochastic neighbor embedding (tSNE). The numbers represent the numbers of learning trials.

## 3. Discussion

Statistical learning is an innate function and starts at an early age for language and music learning regardless of its culture. This study examined how children’s songs are acquired through statistical learning, using the computational model mimicking the statistical learning processes of the brains. We conducted a simulation experiment to visualize the temporal dynamics of perception and production processes through statistical learning among different cultures. The results suggested that, through statistical learning, the models gradually reduced Bayesian surprise that occurred by learning, increased the number of chunks and hierarchy of knowledge. Furthermore, in production, statistical learning leads to the gradual increase of delta-band rhythm. Finally, through statistical learning, new music can be generated gradually in comparison to an original song that was learned. These findings were observed consistently, irrespective of culture. These findings indicate that the innate statistical learning capacity of the brain contributes to the acquisition and generation of delta-band rhythm, chunking, and hierarchy, irrespective of cultural background.

### 3.1. Statistical Learning to Phase Entrainment

The hierarchical structure of rhythms is essential for the acquisition of auditory knowledge such as music and language [18]. Particularly, the higher hierarchy corresponds to a frequency band (approximately, 3Hz>) of prosody, intonation, and long musical note (e.g., half notes), while the lower hierarchy corresponds to a frequency band (approximately, 3Hz<) of syllable and musical notes (e.g., quarter notes) [19]. Such a hierarchical structure of rhythm can be visualized from the AM envelope of sound waveforms [20,21]. Evidence has shown that neural oscillation is entrained in the AM hierarchy of auditory rhythm [22]. Such a phase entrainment contributes to the parsing of the sound signal into each unit of the rhythm hierarchy [23].

The findings in this study demonstrated that statistical learning plays an important role in the acquisition of slower rhythms (<3Hz) regardless of music-cultural differences. A recent study by Attaheri et al. (2022) [26] has highlighted the importance of neural processing of the slower rhythm, specifically, the oscillatory phase entrainment of the delta-band rhythm, for early auditory learning. Previous studies have also demonstrated that the individual capacity for phase entrainment at the lower-frequency band is associated with their statistical learning ability [27]. Furthermore, the neural oscillations appear to synchronize with a statistically chunked rhythm acquired through statistical learning [28,29]. These findings suggest that statistical learning processes play a critical role in the acquisition and consolidation of the slower rhythm.

Our findings also indicated that as the slower rhythms gradually increased, the faster rhythms gradually decreased, highlighting the relationships between these rhythms. It has been suggested that the oscillators are dynamically modulated from slower to faster bands in a top-down manner [22,36]: delta oscillators modulate the theta oscillators, and theta oscillators modulate the gamma oscillators. Such a ‘cascade’ oscillatory system is thought to contribute to encoding the rhythm hierarchy and thus parsing of large (e.g., prosody or phrase) and smaller (e.g., syllables or tone) units in a top-down manner [22].

Auditory information directed towards children, such as speech directed towards infants and children, has been found to have a stronger rhythm in the delta range and a weaker rhythm in the theta range compared to auditory information directed towards adults [37]. Further, another study comparing the hierarchical rhythmic structure across various types of sounds (nature, speech, instrumental music, song, animal sounds, etc.) has demonstrated that the slower band hierarchy (<3 Hz) is especially pronounced in children’s songs and infant/child-directed speech compared to other types of sounds [19] and possibly regardless of culture [25]. Thus, auditory stimuli directed towards children commonly exhibit a stronger power of slow rhythms. Evidence has shown that infant is first phase-entrained to the slower rhythm [26], suggesting that such a slower-band rhythm is important for early learning of language and music. Thus, it is possible that child-directed speech and songs may emphasize the slower rhythms because they syntactically and acoustically learn these rhythms first.

### 3.2. Prediction Error and Chunking

As the statistical learning progressed, it became evident that Bayesian surprise decreased across all songs and cultures. It has been known that the predictive processing of music works to minimize the prediction error between the bottom-up auditory signals and the top-down predictive signals based on internal music models acquired through statistical learning [3,4]. The findings in the present study may imply that statistical learning successfully worked to minimize the prediction error as well as generate chunks, which is a core function in statistical learning.

This study also detected both the increase in the number of chunks and hierarchy as the statistical learning progressed. Researchers have proposed two types of hierarchical statistical learning systems [10,16]. The first system constitutes the fundamental function of statistical learning, which groups chunks of information with high transition probabilities and integrates them into a cohesive unit. The second system involves statistical learning that arranges various chunked units to form a hierarchical syntactic structure (Figure 1). The finding in this study may suggest that the “iteration” of statistical learning is necessary to produce many chunks, which leads to hierarchically structured building.

The t-SNE analysis indicated that, through statistical learning, new music can be generated gradually in comparison to an original song that was learned. This suggests that statistical learning contributes to the ability to generate novel music. An individual internal model (probability distribution) that differs from the probability distribution of the learned song can be constructed by chunking through statistical learning. Such an internal model has a hierarchical structure depending on the amount of learning. This hierarchy contributes to combining between different chunks that was not present in the original song. This combining may facilitate the generation of new music.

Figure 1 shows an example of such a process based on the hierarchical statistical learning of music. Music has general regularities in chord sequences such as the so-called “two-five-one (II–V–I) progression.” Such syntactic progression frequently occurs in music, and therefore, the statistics of the sequential information have high transitional probability and low uncertainty. Thus, once a person has learned the statistical characteristics, it can be chunked as a commonly used unit among musicians. In contrast, the ways of combining the chunked units (e.g., combining among different blue circled units) are different between musicians, leading to the generation of novel music. It is possible that such integration of different chunked units induces human creativity. Further studies are needed to fully understand the relationship between statistical learning and creativity. The present study’s findings may provide important insights to clarify this relationship.

Neurophysiological and behavioral studies have mostly examined differences in cognitive basis across music culture and experiences resulting from statistical learning [12,13]. However, it remained unknown about the temporal dynamics of the emergence of cultural universal systems during the learning process. Using a computational model, this study could gain a constructive understanding of such temporal dynamics including learning and production across different musical cultures.

Our findings suggested that cultural differences may not significantly modulate the statistical learning processes since statistical learning is essential function in the human brain across cultures. Furthermore, statistical learning of children’s songs leads to the acquisition of hierarchical knowledge and the ability to generate novel music. This study provides a novel perspective on the developmental origins of creativity and the importance of statistical learning through early development.

## 4. Conclusion

The present study provided a comprehensive understanding of the role of statistical learning in the acquisition of hierarchy, chunking and slower rhythm, which is critical for acquiring language and music. It is suggested that music-cultural differences may not significantly modulate the statistical learning processes. Furthermore, statistical learning may lead to the generation of novel music. These findings have important implications for our understanding of neural processing and cognitive development, particularly in the context of auditory learning.

## 5. Materials and Methods

### 5.1. Hierarchical Bayesian Statistical Learning Model

This study developed a computational model, which simulates statistical learning processes of the brain, referred to as HSBL model [30]. The scripts of the model have been deposited to an external source (https://osf.io/cqmz8/?view_only=b95d94626a364700adb9e1e94384525d). This is a model that integrates Bayesian estimation with Markov processes using a Dirichlet distribution as a prior distribution. This model can not only calculate the transition probabilities but also determine the “reliability of the transition probabilities” from the inverse of the variance of the prior distribution of the transition probabilities. Using the normalized values of transition probabilities and reliability, this model chunks transition patterns when the product of “reliability * probability” is greater than a constant c. The constant can be decided based on the sample length and the number of learning trial. In this study, we defided c=5 given the mean sample length used in this experiment. A chunked unit can be further integrated with another chunked unit, leading to generation of a longer unit in higher hierarchy (see Figure 1). That is, by the cascade of chunking during statistical learning, the model gradually forms the hierarchical structure.

### 5.2. Materials and Learning and Production Processes

This study generated 15 different models by manipulating the amount of learning. We used the MIDI data of 364 children’s song including 50 English, 80 German, 80 Spanish, 74 Japanese, and 80 Korean songs as the training data, and repeated the learning of the song one to fifteen times in each of 364 songs (the titles of music were shown in the supplementary material). We investigate how the model transforms the internal model over 15 trials of learning. Furthermore, using the probability distribution of these 15 models, 20 pieces were probabilistically generated for each model through an automatic composition process [30].

### 5.3. Comparison of Internal Representations in the Model

This study compared the total Bayesian surprise (or total prediction errors) that occurred during learning, measured by the Kullback-Leibler divergence between a distribution P(x) before learning an event (e_n_) and a distribution Q(x) after learning the event (e_n+1_), as well as the total number of chunks generated during 15 trials of statistical learning. The Kullback-Leibler divergence has often been used to measure prediction error or Bayesian surprise in the framework of predictive processing of the brain [2,31,32]. It is a metric used to measure the similarity between two different probability distributions. It represents how much information is lost when one probability distribution changes into another, and since it is non-negative, a small value indicates that the two distributions are similar. Specifically, it is calculated by taking the difference between the probability density functions of the two distributions, taking the logarithm at each point, and then computing the weighted average with respect to one of the distributions. The Kullback-Leibler divergence between two probability distributions P(x) and Q(x) is calculated using the following formula:

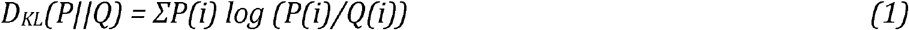

Here, P(i) and Q(i) represent the probabilities of selecting the value i according to the probability distributions P and Q, respectively. The number of chunks and hierarchy, and values of Bayesian Surprise were normalized to z score. In addition, we calculated the average probability distribution of the 20 songs generated by each model and compared the similarity of the models to the training data (i.e., original data) using t-distributed stochastic neighbor embedding (tSNE).

### 5.4. Comparison of Acoustic Properties of Rhythm

We converted the MIDI data of the 20 songs generated by each model into WAV format and extracted the rhythm waveform (modulation wave) below 15 Hz using the Bayesian probabilistic amplitude demodulation model (PAD, [20]). The acoustic signals were first normalised based on the z-score (mean = 0, SD = 1) in case the sound intensity influenced the spectrotemporal modulation feature. The spectrotemporal modulation of the signals was analysed using PAD to derive the dominant AM patterns. music and speech signals can be decomposed into slow-varying AM patterns and rapidly-varying carrier or frequency modulation (FM) patterns [19,33-35]. AM patterns are responsible for fluctuations in sound intensity, which are considered to be a primary acoustic feature of perceived hierarchical rhythm. On the other hand, FM patterns reflect fluctuations in spectral frequency and noise. One can separate the AM envelopes of speech signals from the FM structure by means of amplitude demodulation processes. The PAD model employs Bayesian inference to infer the modulators and carrier, and to identify the envelope that best fits the data and a priori assumptions. More specifically, amplitude demodulation is the process by which a signal (yt) is decomposed into a slowly varying modulator (mt) and a rapidly varying carrier (ct):

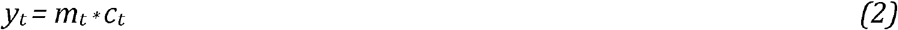

PAD employs amplitude demodulation as a process of both learning and inference. Learning involves the estimation of parameters that describe distributional constraints, such as the expected timescale of variation of the modulator. Inference involves estimating the modulator and carrier from the signals based on learned or manually defined parametric distributional constraints. This information is probabilistically encoded in the likelihood function *P(y*_*1:T*_|*c*_*1:T*_, *m*_*1:T*_, *θ)*, the prior distribution over the carrier *p(c*_*1:T*_|*θ)*, and the prior distribution over the modulators: *p(m*_*1:T*_|*θ)*. Here, the notation x_1:T_ represents all the samples of the signal x, ranging from 1 to a maximum value T. Each of these distributions depends on a set of parameters θ, which control factors such as the typical timescale of variation of the modulator or the frequency content of the carrier. For more detail, the parametrised joint probability of the signal, carrier, and modulator is:

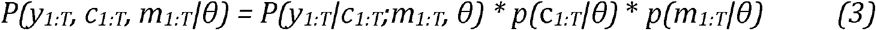

Bayes’ theorem is applied for inference, forming the posterior distribution over the modulators and carriers, given the signal:

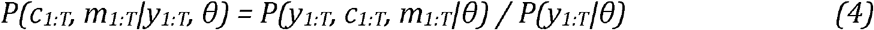

The full solution to PAD is a distribution over the possible pairs of modulators and carriers. The most probable pair of modulator and carrier given the signal is returned:

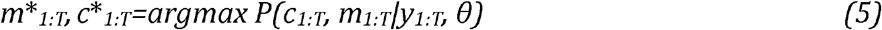

PAD utilizes Bayesian inference to estimate the most suitable modulator (i.e., envelope) and carrier that best align with the data and a priori assumptions. The resulting solution takes the form of a probability distribution, which describes the likelihood of a specific setting of modulator and carrier given the observed signal. Thus, PAD summarizes the posterior distribution by returning the specific envelope and carrier with the highest posterior probability, thereby providing the best fit to the data.

PAD can be run recursively using different demodulation parameters each time, producing a cascade of amplitude modulators at different oscillatory rates to form an AM. The positive slow envelope is modeled by applying an exponential nonlinear function to a stationary Gaussian process, resulting in a positive-valued envelope with a constant mean over time. The degree of correlation between points in the envelope can be constrained by the timescale parameters of variation of the modulator (i.e., envelope), which can either be manually entered or learned from the data.

In the present study, we manually entered the PAD parameters to produce the modulators at an oscillatory band level (i.e., <10 Hz) isolated from a carrier at a higher frequency rate (>10 Hz). The carrier reflects components, including noise and pitches, for which the frequencies are much higher than those of the core modulation bands. In each sample, the modulators (envelopes) were converted into time-frequency domains using scalogram. The scalograms depict the AM envelopes derived by recursive application of probabilistic amplitude demodulation. We then calculated the average frequency power at each frequency and further averaged it over the 20 songs generated by each model.

## Acknowledgements

This research was supported by JSPS KAKENHI (22KK0157; 22H05210; 21H05063) and JST Moonshot Goal 9 (JPMJMS2296), Japan. The funding sources had no role in the decision to publish or prepare the manuscript.

## Supporting information captions

Information of Music Corpus

